# Analysis of natural structures and chemical mapping data reveals local stability compensation in RNA

**DOI:** 10.1101/2024.12.11.627843

**Authors:** Robert L. Cornwell-Arquitt, Riley Nigh, Michael T. Hathaway, Joseph D. Yesselman, David A. Hendrix

**Author notes:** Correspondence should be addressed to David A. Hendrix and Joseph D. Yesselman.

## Abstract

RNA molecules adopt complex structures that perform essential biological functions across all forms of life, making them promising candidates for therapeutic applications. However, our ability to design new RNA structures remains limited by an incomplete understanding of their folding principles. While global metrics such as the minimum free energy are widely used, they are at odds with naturally occurring structures and incompatible with established design rules. Here, we introduce local stability compensation (LSC), a principle that RNA folding is governed by the local balance between destabilizing loops and their stabilizing adjacent stems, challenging the focus on global energetic optimization. Analysis of over 100,000 RNA structures revealed that LSC signatures are particularly pronounced in bulges and their adjacent stems, with distinct patterns across different RNA families that align with their biological functions. To validate LSC experimentally, we systematically analyzed thousands of RNA variants using DMS chemical mapping. Our results demonstrate that stem folding, as measured by reactivity, correlates with LSC (R^2^ = 0.458 for hairpin loops) and that instabilities show no significant effect on folding for distal stems. These findings demonstrate that LSC can be a guiding principle for understanding RNA function and for the rational design of custom RNAs.

**Graphical Abstract:** 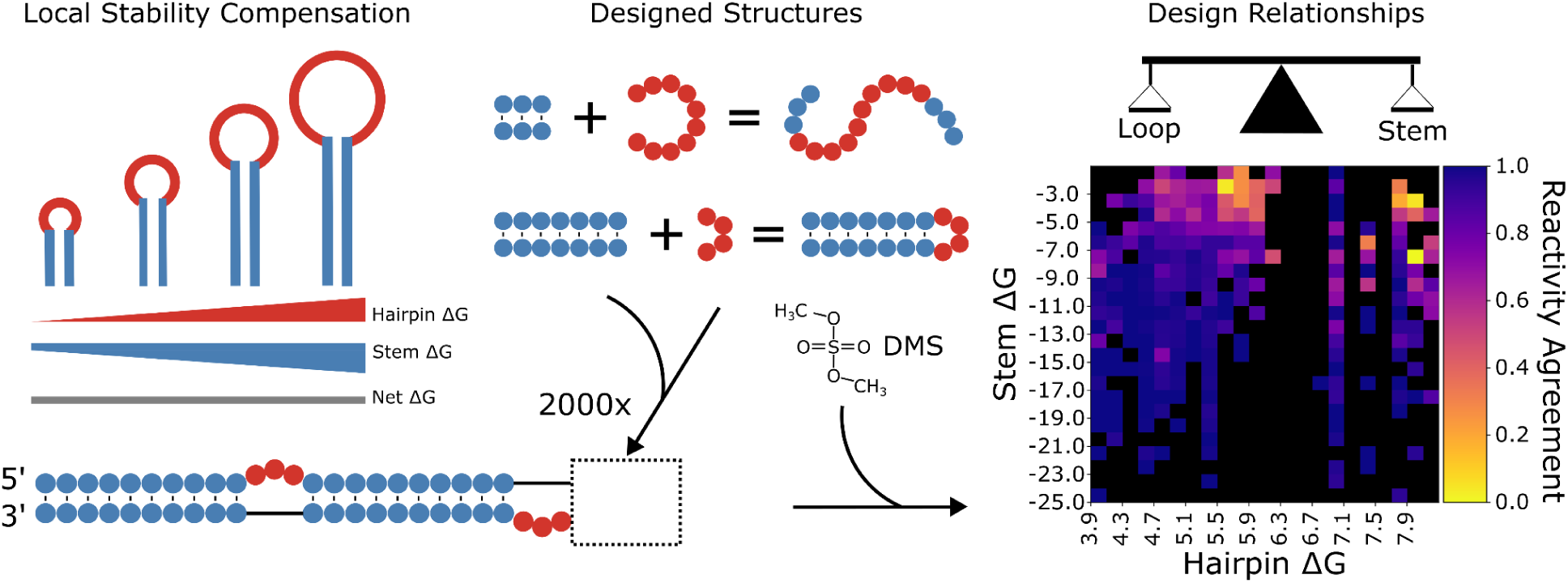

## Introduction

RNA molecules adopt complex structures that perform essential functions across biology, from regulating gene expression to modifying chromatin structure (1–3). This structural versatility has inspired bioengineers to harness RNA as a platform for creating synthetic molecular machines, including scaffolds that colocalize enzymes (4–6), custom-designed ribozymes (7–9), and aptamers with specific ligand-binding properties (10–12). The ability of RNA to perform these varied functions fundamentally depends on its ability to fold into three-dimensional (3D) structures. Therefore, advancing our understanding of RNA folding mechanisms is crucial for elucidating the behavior of biological RNAs and developing more effective RNA-based therapeutics and engineering applications.

The field utilizes diverse experimental approaches to understand RNA folding, ranging from high-resolution techniques like X-ray crystallography (13–15) and cryo-electron microscopy (16–18) to lower-resolution methods like chemical mapping (19–21) and cross-linking (22–24). Yet fundamental challenges remain in the prediction of an RNA’s base-pairing pattern (secondary structure) from sequence alone, which would reduce the need for time-consuming experimental characterization of each new RNA (25). Significant progress in computational prediction began with the development of the "Turner parameters," which formed the foundation for algorithms like Mfold (26), RNAStructure (27), RNAfold (28) that calculate the global minimum free energy (MFE) structure. The field continues to evolve rapidly, with linear-time structure prediction methods (29), machine learning (30), and deep learning approaches enhancing our ability to predict RNA secondary structures with increasing accuracy (31–33), but they often struggle when presented with novel RNAs (34) or RNAs from families that are different from what they were trained on. Therefore, the identification of new patterns that could improve RNA structure prediction remains highly valuable.

Structure prediction algorithms can also be leveraged to identify sequences that will fold into a desired secondary structure for engineered applications. This “inverse-folding” problem is computationally intensive due to the vast sequence space available for any base-pairing arrangement. Current approaches often focus on finding sequences that optimize the minimum free energy (MFE) (28, 35, 36). However, these strategies result in sequences with high GC content, which often misfold (37). The limitations of high-GC designs have been demonstrated empirically through the collective wisdom of Eterna participants, who found that excessive GC content diminishes the success of RNA design tasks across thousands of experiments (38). Notably, nature does not employ this strategy — ribosomal and transfer RNAs maintain consistent GC content despite variation in background GC content (39). This observation suggests that natural RNA sequences follow alternative optimization principles that could inform more effective approaches to designing synthetic RNA structures.

The MFE approach considers the global sum of energy terms, and fails to account for the spatial distribution of stability within an RNA structure. For example, larger RNA loops have been hypothesized to require larger stabilizing helices to offset the entropic cost of loop formation (40). By simply minimizing free energy globally, this energetic requirement may be satisfied by helices located anywhere in the structure. Nature may allocate its limited free energy "budget" according to functional priorities and to specific locations in need of greater stabilization rather than global minimization. We propose that the local energetic balance between loops and their adjacent stems—a relationship we term local stability compensation (LSC)—may be a design rule that nature employs and could be leveraged in rational RNA design.

We propose that larger and more destabilizing loops in structures exhibiting LSC must be paired with proportionally larger and more stabilizing stems to ensure consistent folding for functional RNAs that rely on robust structures. However, LSC may not be strictly necessary for RNAs that lack evolutionary pressure for robust folding or for dynamic RNA substructures that adopt many conformations. Therefore, apart from such examples, we expect to observe a pattern where decreasing stem ΔG balances increasing loop ΔG to maintain a consistently negative net free energy (ΔG) (**Supplementary Fig. 1A**). Crucially, this compensation operates locally: each loop is primarily stabilized by its adjacent stems, with minimal influence on distant structural elements.

In this study, we first analyzed LSC signatures in natural RNA structures, revealing distinctive patterns in stem and loop ΔG distribution across different loop types and RNA classes. We then experimentally tested the relationship between LSC and its impact on folding by performing DMS chemical mapping on a library of designed RNAs. Our results demonstrate that folding fidelity correlates with LSC and confirm that structural instabilities primarily affect local folding. These findings have significant implications for RNA design strategies, suggesting that meeting local stability requirements is crucial for successful substructure folding. Furthermore, understanding the stability requirements of individual substructures could enhance RNA design by facilitating their assembly into larger, more complex structures.

## Materials and Methods

### Analysis of net ΔG for local substructures with bpRNA-1m

For our large scale analysis of RNA structures, we used the meta-database bpRNA-1m (41). A smaller subset without structures of 90% or greater sequence identity, called bpRNA-1m90 was also used to avoid over sampling particular RNAs. Importantly, structural data in the database are accessible as structure type (.st) files which contain a breakdown of the subcomponents of the RNA structure to facilitate our analysis. We calculated the free energies of the individual substructures using the Turner 2004 parameters for RNA folding (42). We define **net ΔG** as the sum of the stabilizing stem free energy and the destabilizing adjacent loop free energy. In the case of hairpin loops, there is only one stem/helix immediately adjacent to the loop. For two-way junctions (i.e. bulges and internal loops), we average the free energies of both stems on each side of the junction.

We developed a pipeline using Vienna RNA to refold structures from the RNA families database (RFAM) and the comparative RNA web (CRW) database using the consensus base pairs as hard constraints in order to correct for “under folded” structures which contain large stretches of unpaired nucleotides. The resulting dataset was used for the analysis of large-scale trends across many RNAs (**Fig. 1**), and for correcting unrealistic loops in particular RNAs of interest (**Fig. 3B-D**). The dataset was not used when the numbering scheme of conserved loops needed to be preserved.

**Figure 1.**
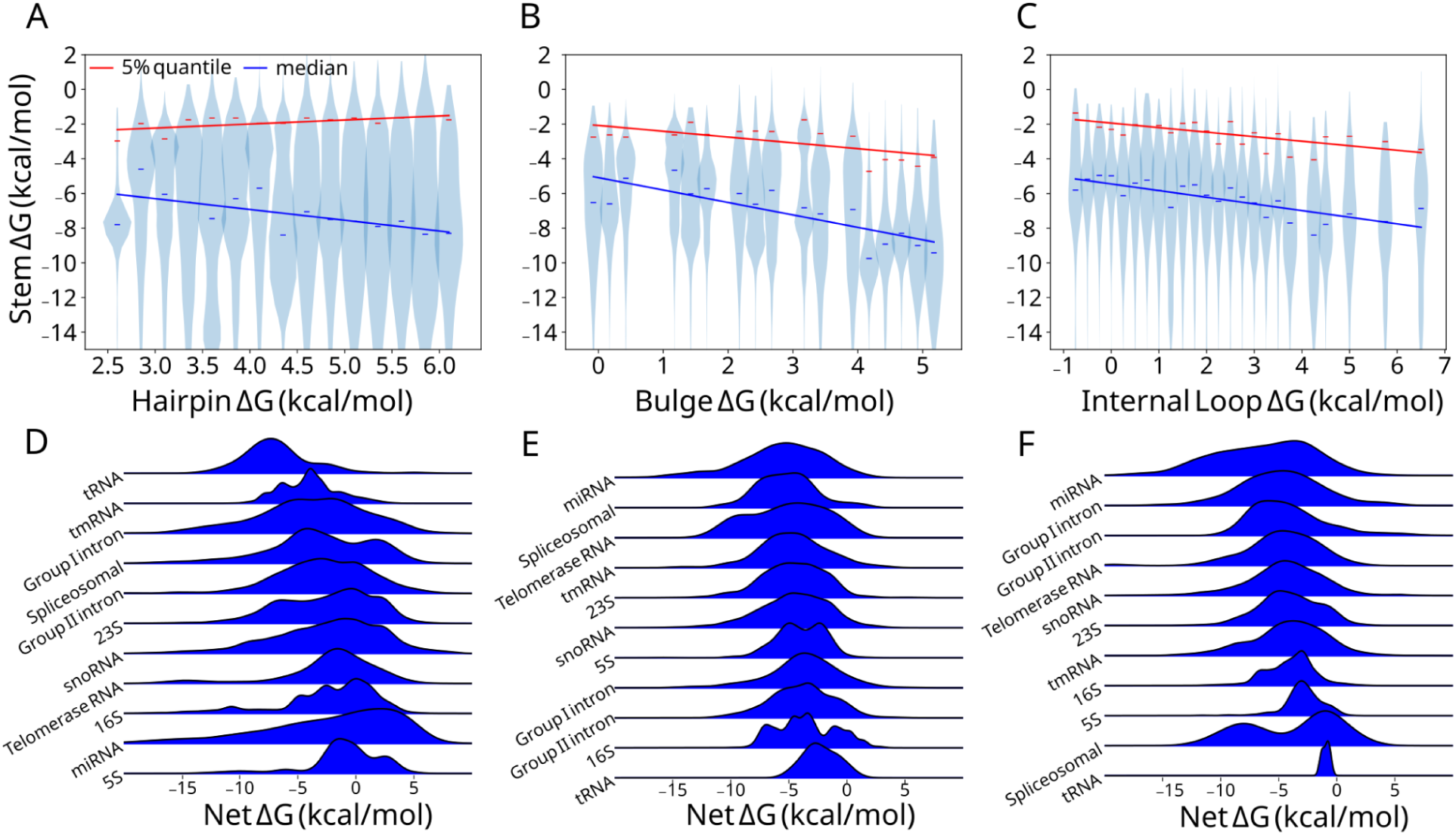
Local stability compensation in bpRNA-1m90. (A) Hairpin, (B) bulge, and (C) internal loop ΔGs from refolded bpRNA-1m90 structures are binned and the corresponding stem ΔGs are plotted as violins with linear regressions drawn for the median (blue) and 5% quantile (red). (D) bpRNA-1m90 net free energy kernel density estimates for hairpin loops and (E) bulges and (F) internal loops belonging to different RNA types.

### Data analysis and statistics

All statistics were performed using python code (https://doi.org/10.5281/zenodo.14252189) with the SciPy and Scikit-learn modules. For assessing the significance of net ΔG distributions in hairpin loops, bulges, and internal loops in Supplementary Figure 1B, an in-structure rotation control was used: loop energies were summed with stem energies belonging to a distal substructure instead of loop-adjacent stem energies. This was only done for structures where more than three loops of a type existed to eliminate the possibility of a loop receiving one or more of its own stems. The difference over control in net free energy frequencies by bin was calculated along with an f test to reveal the significance of differences in variance.

392 hairpin sequences were filtered out from the dimethyl sulfate (DMS) reactivity results because of C repeats greater than 4 residues which may result in unreliable DMS reactivities as seen in A repeats (43). DMS data for 875 hairpin loops, 1979 bulges, and 1987 internal loops was available for analysis.

For analysis of DMS reactivity data, we employed the area under the receiver-operator curve (AUROC) to quantify folding fidelity, that is, the agreement of reactivity values with the ‘designed’ structure. This measure quantifies the overlap between the reactivity distributions of paired-by-design and unpaired-by-design nucleotides for a single RNA structure. In this framework, unpaired nucleotides are treated as the “positive predictions” and paired nucleotides are treated as the “negative predictions” for the AUROC calculation, while reactivity data provides the score to be compared. We calculated the AUROC using the Scikit-learn python package for the AUROC score across adenine/cytosine (A/C) nucleotides in global, local, and distal scopes (all residues, residues in the residues region, and residues not in the randomized region, respectively). In addition, we measured the average reactivity of stems and loops belonging to the randomized region and a region distal to the randomized region as a control. Only reactivities corresponding to A or C nucleotides were used for these calculations. The sample sizes depended on the size of the substructure and the abundance of A/C residues (see library design for more details). Lastly, a set of per-position averages were calculated with 95% confidence for various bins of net ΔG, which involved sample sizes no lower than 70 nucleotides for each bin.

### Substructure free energy calculation and designed RNA library generation

We developed a python module to calculate the free energies of folding by components called bpRNAStructure. We used bpRNAStructure to extract tab-delimited files containing bpRNA-1m data organized by RNA ID number and the substructure identifier. These data were used in the aforementioned analyses and statistics regarding bpRNA-1m. We also used bpRNAStructure to add the free energy component-wise as an additional annotation of the structure type file from bpRNA-1m, hence calling it a “ste” file for “structure type energy”. These files were used to organize data for each sequence in the designed RNA libraries, which are introduced below.

We used a python script to randomly generate sequence libraries with an important set of constraints. Each sequence was based on a core template and required: a hamming distance of at least 20 compared to any other sequence, loops of A and C residues, and substructure lengths of 4-12, 3-11, 1-9, and 1-9 residues for stems, hairpin loops, bulges, and internal loops respectively. The template was a triply bulged hairpin structure, with variable and semi-constant regions depending on the loop type being studied (**Fig. 4A**). The variable regions consisted of loops varying by length and stems varying by length, GC content, and G:U pairs. ‘Constant’ energy stems were used for non-varying (distal) regions and were designed to minimize free energy variation while allowing for sequence variation. Constant energy stems used G:C or C:G closing base pairs, a fixed GC content, and 0 G:U pairs. Because sequences were required to be no shorter or longer than 20% of the mean sequence length, 21 and 13 sequences were filtered out of the internal loop and bulge libraries respectively. Oligonucleotide pools of each library were ordered from Agilent Technologies.

### PCR amplification of DNA templates

DNA templates for transcription were prepared by PCR amplification of the synthesized oligonucleotide pool. We first resuspended the oligo pool in 50 μL of IDTE buffer (1X, pH 8.0, IDT #11-05-01-13). PCR amplification was performed using custom primers obtained from IDT: a forward primer containing the T7 promoter sequence (TTCTAATACGACTCACTATAGG) and a reverse primer (GTTGTTGTTGTTGTTTCTTT). Each 50 μL PCR reaction mixture contained Q5 High-Fidelity DNA Polymerase (25 μL, NEB #M0494S), oligo pool template (2 μL), forward and reverse primers (2.5 μL each at 10 μM, diluted from 100 μM stocks), and RNase-free UltraPure water (18 μL, ThermoFisher #10977015). The PCR thermal cycling conditions consisted of initial denaturation at 98°C for 30 seconds, followed by 20 cycles of: denaturation at 98°C for 10 seconds, annealing at 62°C for 15 seconds, and extension at 72°C for 15 seconds. A final extension step was performed at 72°C for 5 minutes. The PCR products were resolved by electrophoresis on a 2% agarose gel (150V for 1 hour) and extracted using the Zymoclean Gel DNA Recovery Kit (Genesee Scientific #11-301C).

#### *In vitro* transcription of RNA libraries

For *in vitro* transcription of the RNA libraries, we first prepared: 10x Transcription (Tx) Buffer containing 400 mM tris-base, 10 mM spermidine, and 0.1% Triton X; 25 mM NTPs, 50 mM DTT, and 250 mM MgCl_2_. The transcription reaction contained: 10 μL of 10x Tx Buffer, 5 μL of 50 mM DTT, 16 μL of 25 mM NTP solution, 8 μL of 250 mM MgCl_2_, 4 μL of T7 RNA polymerase (New England Biolabs #M0251S), and 33 μL RNase-Free water. Purified template DNA was quantified using a NanoDrop spectrophotometer, adjusted to 0.3 μM, and 24 μL was added to complete the reaction mixture. The complete reaction was incubated at 37°C for 6 hours. Following transcription, the template DNA was removed by DNase I digestion using the RNA Clean and Concentrator-5 with DNase I kit (Genesee Scientific #R1014). The RNA product was purified using the same kit according to the manufacturer’s instructions. Prior to DMS MaPseq analysis, we performed quality control by measuring RNA concentration by NanoDrop spectrophotometer and confirming the RNA length by electrophoresis on a 4% denaturing agarose gel run at 150 volts for 1 hour.

### DMS probing of RNA libraries

For each RNA library, we prepared a solution containing 10 pmol of RNA in 5 μL RNase-Free water. RNA samples were denatured by heating to 90°C for 4 minutes followed by rapid cooling to 4°C for 3 minutes in a thermocycler. The denatured RNA was then added to a folding buffer containing 16.5 μL 0.4 M sodium cacodylate buffer and 1 μL of 250 mM MgCl_2_, yielding final concentrations of 264 mM sodium cacodylate and 10 mM MgCl_2_ during the DMS reaction. The RNAs were folded at room temperature for 30 minutes. A fresh DMS solution was prepared by combining 15 μL DMS (Sigma-Aldrich #D186309) with 85 μL 100% ethanol (Decon Labs cat. #2716). Immediately following the 30-minute folding period, 2.5 μL of the freshly prepared DMS solution was added to the RNA-buffer mixture to initiate modification. The reaction was allowed to proceed for 6 minutes before being quenched with 25 μL BME (ThermoFisher cat. #125470010). Modified RNA was purified using the RNA Clean & Concentrator-5 kit (Genesee Scientific #R1014) and eluted in 7 μL of RNase-Free water. Final RNA concentration was determined using the Qubit RNA BR Assay Kit (ThermoFisher #Q10211), using 1 μL of the purified RNA sample for measurement.

To detect adenine and cytosine methylation sites, we employed Marathon reverse transcriptase (Kerafast #EYU007), which incorporates mutations during cDNA synthesis at methylated positions. Prior to reverse transcription, we prepared the following solutions: 5x Marathon buffer (250 mM Tris-HCl pH 8.3, 375 mM KCl, 15 mM MgCl_2_), 10 mM dNTPs, 100 mM DTT. The modified RNA sample was diluted to 0.25 μM, and the RT primer (**Supplemental Document: Sequences.xlsx**) was diluted to 0.285 μM. The reverse transcription reaction was assembled by combining 3 μL of 5x TGIRT buffer, 1.49 μL of 10 mM dNTPs, 0.74 μL of 100 mM DTT, 1.2 μL of Marathon reverse transcriptase, 6.4 μL of diluted modified RNA, and 1 μL of diluted RTB primer. The reaction was thoroughly mixed and incubated at 42°C for 3 hours. Following incubation, we added 5 μL of 0.4 M NaOH to the reaction and subjected the mixture to a denaturation step (90°C for 4 minutes followed by cooling at 4°C for 3 minutes). The NaOH was then neutralized by adding 2.02 μL of Quench Acid (1.43 M NaCl, 0.57 M HCl, 1.29 M sodium acetate). To purify the cDNA product, we first added 27.5 μL of RNase-Free water to the neutralized reaction, then purified using the Oligo Clean and Concentrator Kit (Genesee Scientific #11-380B). The final cDNA product was eluted in 15 μL of RNase-free water.

The cDNA products were amplified by PCR using primers obtained from IDT (dissolved in IDTE pH 8 buffer at 100 μM). We used the forward primer AATGATACGGCGACCACCGAGATCTACACTCTTTCCCTACACGACGCTCTTCCG and the reverse primer CAAGCAGAAGACGGCATACGAGATCGGTCTCGGCATTCCTGCTGAACCGCTCTTCCGATCTGGTACTATGTACCAAAG. Prior to PCR setup, primers were diluted 1:10 to 10 μM in RNase-free water.The PCR reaction contained 25 μL Q5 High-Fidelity DNA Polymerase (New England Biolabs #M0494S), 5 μL each of diluted forward and reverse primers, 6 μL purified cDNA, and 9 μL RNase-free water. Thermal cycling conditions consisted of an initial denaturation at 98°C for 30 seconds, followed by 16 cycles of: denaturation at 98°C for 10 seconds, annealing at 62°C for 15 seconds, and extension at 72°C for 15 seconds. A final extension step was performed at 72°C for 5 minutes. For gel purification, we first prepared a 2% agarose gel by diluting 10x TBE Buffer (Bio-Rad #1610770) to 1x with RNase-free water, then dissolving 1.5g agarose (Apex #20-102) in 75 mL of 1x TBE buffer. The solution was heated in a microwave for 2 minutes, mixed thoroughly, and supplemented with 10 μL SYBR Safe DNA Gel Stain (ThermoFisher #S33102) before casting. The Fisherbrand Horizontal Electrophoresis System (Fisherbrand #FBSB710) was used for electrophoresis, with 1x TBE as a running buffer. PCR products were mixed with 10 μL TackIt Cyan/Yellow Loading Buffer (ThermoFisher #10482035) and separated on the gel at 150 volts for 60 minutes. After imaging, bands of the correct size were excised and purified using the Zymoclean Gel DNA Recovery Kit (Genesee Scientific #11-301C). Final DNA concentration was determined using the Qubit 1X dsDNA High Sensitivity Assay Kit (ThermoFisher #Q33230).

### DMS-MaPseq data analysis

Sequencing was performed on a Novaseq 6000. Each sequencing run was first demultiplexed using the RTB barcodes added during the RT. Demultiplexing was performed using novobarcode (https://www.novocraft.com/documentation/novobarcode/demultiplexing-barcodedindexed-reads-with-novobarcode/).

novobarcode -b rtb_barcodes.fa -f test_R1_001.fastq test_R2_001.fastq

where rtb_barcodes.fa gives a list of sequencing barcodes, such as

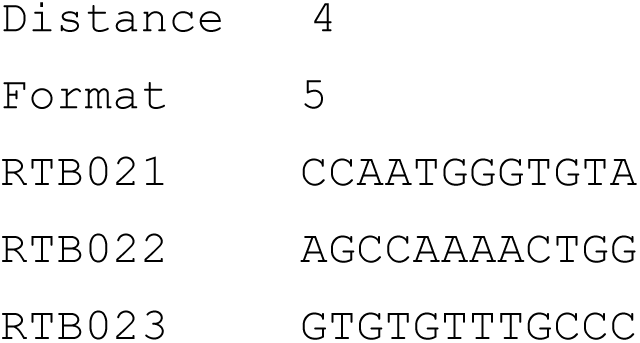

Where three barcodes are specified, distance is the distance in base pairs between a barcode and an allowable read. Format specifies that the barcode will be on the 5′ end of the read 1.

Lastly, we used the RNA mutational profiling (RNA-MaP) tool to count mutations to determine the mutational fraction at each nucleotide position (https://github.com/YesselmanLab/rna_map). This is an open-source tool developed to simplify mutational profiling analysis.

rna-map -fa test.fasta -fq1 test_mate1.fastq -fq2 test_mate2.fastq --dot-bracket test.csv --param-preset barcoded-libraries

The rna-map command requires a FASTA-formatted file containing all DNA reference sequences and the paired sequencing reads generated from the previous step. We also supplied a CSV file containing the dot bracket structure for each RNA with the --dot-bracket flag and applied stricter constraints to how well each sequence needs to align to a read --param-preset barcoded-libraries. All the data are stored in JSON format in a pandas DataFrame.

## Results

### Evidence for local stability compensation found in a structural meta-database

LSC proposes that RNA folding stability requires larger, destabilizing loops to be paired with more stabilizing adjacent stems (Supplementary Fig. 1A). To test whether LSC is observed in biological RNAs, we analyzed the relationship between loop and stem free energies (ΔG) across 79,582 hairpins, 30,441 bulges, and 17,117 internal loops in the bpRNA-1m90 database. Importantly, RFAM structures can feature unrealistically large loops originating from nucleotide stretches that do not align with the consensus structure. We applied a pipeline (see methods) to refold these loops holding consensus base pairs fixed resulting in targeted corrections that introduced shifts in the bulk distributions, particularly a depletion of the highest net ΔG loops. (**Supplementary Fig. 2**). In the resulting data, hairpin loop, bulge, and internal loop ΔG bins of 0.25 kcal/mol correlate negatively with the median stem ΔG (**Fig. 1A-C**), with bulges appearing to correlate more strongly (R^2^ = 0.647)(**Supplementary table 1**). In order to assess the upper limit of stem free energy for associated loops, we compared the 5% weakest stem ΔGs (**Fig. 1A-C**). The 5% quantiles for bulges and internal loops correlate with a slope similar to the median, except for hairpins. Furthermore, bulge and internal loop free energy sum (net ΔG) distributions are enriched between -3 and -6 kcal/mol compared to a control in which stems are swapped with distal stems (see methods), which is not observed in hairpins (**Supplementary Fig. 1B**). These data suggest that LSC exists in bpRNA-1m90 structures, but is partially obscured in hairpins, likely by RNA family-specific factors which may contribute to the inconsistent compensation patterns. It is likely not all RNAs have strong structure requirements and we may see stronger effects examining only RNAs in a given family.

When grouped by their respective RNA family, net ΔG comparisons of hairpins (**Fig. 1D**), bulges (**Fig. 1E**), and internal loops (**Fig. 1F**) revealed distinct stability compensation patterns. Hairpin loops showed the highest variability, with a median net ΔG of -2.66 kcal/mol and a standard deviation of 1.74 kcal/mol, which is higher than that of internal loops (1.33 kcal/mol) and bulges (0.75 kcal/mol) (**Supplemental table 2**). A comparison by more specific RNA subclasses (**Supplementary Fig. 3**) corroborated this observation albeit with greater variation (**Supplementary table 2**). Because 1) the variance by RNA type accounts for some portion of the variance unexplained by the global correlation, and 2) the aforementioned variances by type are inversely proportional to the corresponding global correlations, RNA family-specific compensation patterns are in part responsible for deviations from LSC.

### Local stability compensation varies by RNA type and is consistent with function

We hypothesized that LSC may be more prominent in specific RNA families where high stability is needed. Conversely, we speculated that high net ΔG for some families may indicate functions where destabilization is necessary. To investigate in more detail, we highlight four classes with distinct LSC behaviors. The C4 antisense RNA is a noncoding RNA (ncRNA) contained in P1 and P7 bacteriophages (44, 45) (**Fig. 2A**). Hairpin loops within the C4 antisense RNA were selected for further investigation because they have negative and highly conserved net ΔGs. The C4 antisense RNA contains two highly stabilized hairpins. H1 consists of a highly stable hairpin loop and stem while H2 consists of a large hairpin loop and stable GC-rich stem. Both stem-loops achieve highly similar net ΔGs despite the substantial differences in their subcomponents (**Fig. 2B**).

**Figure 2.**
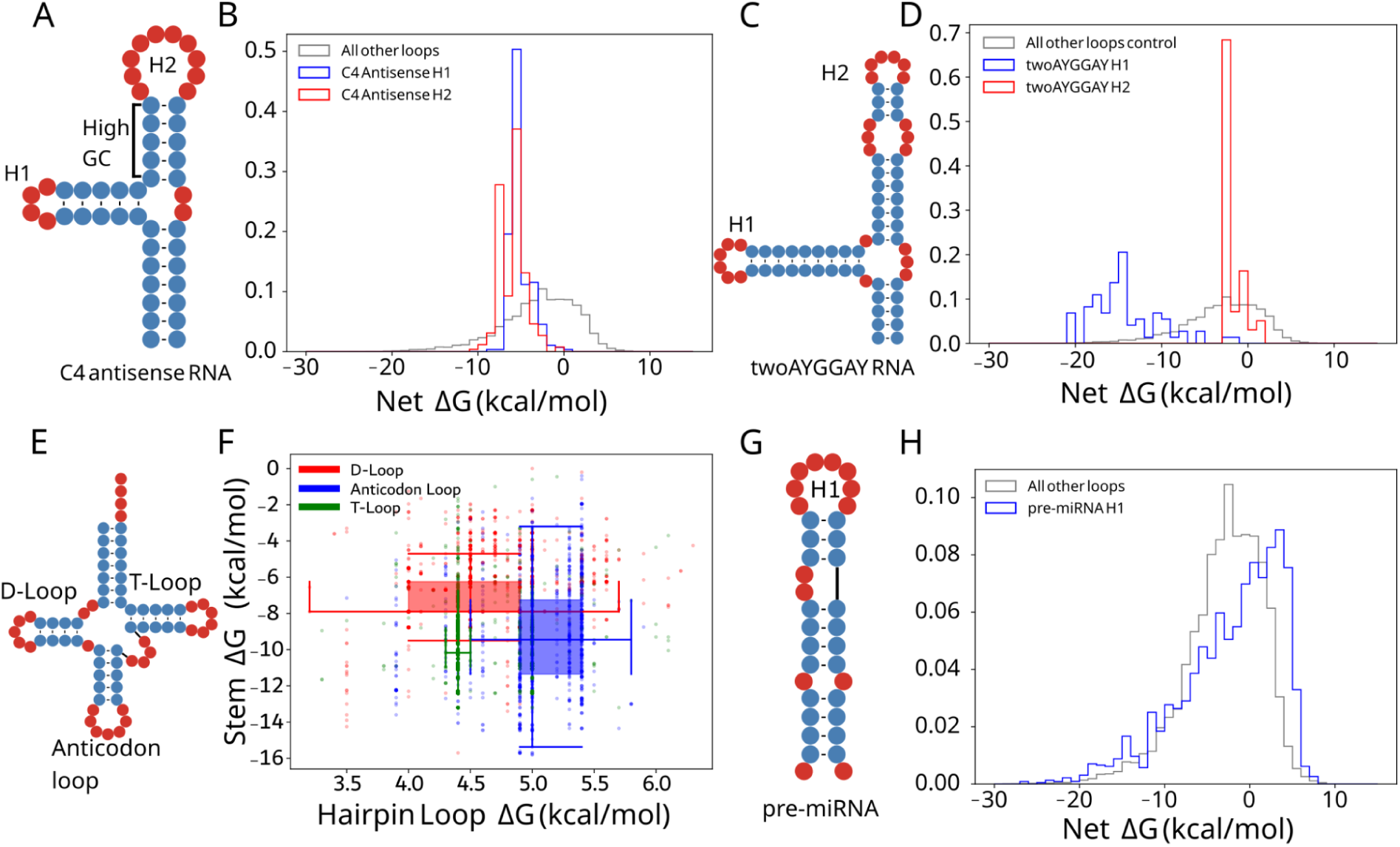
Net ΔG distributions are distinct for RNA types and particular RNA loops. (A) C4 antisense RNA example secondary structure and (B) hairpin net ΔG distribution (H1 in blue, H2 in red) and other hairpins in bpRNA-1m90 (grey). (C) TwoAYGGAY RNA example secondary structure and (D) net ΔG distribution (H1 in blue, H2 in red) and other hairpins in bpRNA-1m90 (grey). (E) Example tRNA secondary structure (bpRNA_RFAM_1424) and (F) two dimensional box plots for the D-loop, T-loop, and anticodon loop in red, green, and blue respectively. (G) pre-miRNA example secondary structure and (H) net ΔG distribution (blue) and other hairpins in bpRNA-1m90 (grey).

The TwoAYGGAY RNA motifs (**Fig. 2C**), characterized by two conserved AYGGAY terminal hairpin loops (45), provided an example of family-specific stability patterns. In a departure from the expectations of LSC and the observations in C4 antisense RNAs, the TwoAYGGAY RNAs showed a bimodal net ΔG distribution, with peaks at -16 and -3 kcal/mol (**Fig. 2D**, **Supplementary Fig. 3A**), revealing that identical hairpin loop sequences can maintain different stabilities. This conserved differential stabilization within the same RNA structure suggests a potential functional significance for this relatively unexplored motif. This RNA class demonstrates that LSC patterns may be of interest in RNAs of unknown function.

Transfer RNAs (tRNAs) are a well-studied class of noncoding RNAs that are involved in translation with a cloverleaf secondary structure containing three hairpins (**Fig. 2E**). We investigated both the stem and loop free energies in tRNAs for energy relationships specific to different hairpins. tRNA hairpins group into three distinct regions of stem and hairpin loop ΔG, shown with box and whisker plots (**Fig. 2F**). While the D-loop and anticodon loop have similar variability, the anticodon loop is larger and higher energy and is also associated with a more stabilized stem. The observation of more destabilizing loops being coupled with more stabilizing stems is consistent with the LSC hypothesis.

MicroRNA (miRNA) precursors (pre-miRNAs) have well-defined structures (**Fig. 2G**) that are conserved for the biogenesis of small RNAs and share a similar function despite substantial sequence and structure variation (46, 47). Indeed, pre-miRNA hairpins in bpRNA-1m90 show a wider net ΔG distribution than all other types (**Fig. 2H**), which may be consistent with the target sequence diversity of miRNAs (47) and the loop positioning important for Dicer function (48). These structures frequently contained poorly stabilized hairpin loops with net ΔGs reaching 5 kcal/mol, far higher than the hairpins typical of other substructures (**Fig. 2H**). This high instability of the stem closing the hairpin loop is notable due to its proximity to the cleavage site. Because of phylogenetic differences in miRNA maturation, we continue this analysis of pre-miRNAs from a phylogenetic perspective.

### Local stability compensation is distinguished by phylogeny

Owing to species-related differences in form and function of miRNAs, and to the thermodynamic nature of LSC, we investigated how LSC correlates with phylogeny. We compared plant and metazoan pre-miRNAs, as well as several RNA types belonging to thermophiles and non-thermophiles. Plant pre-miRNA hairpins were enriched relative to metazoan pre-miRNAs at 5.0 kcal/mol and above (**Fig. 3A**, **Supplementary table 2**), but these structures included many hairpin loops of unrealistic sizes. To account for this, we employed the results of structure prediction with constraints (See methods). In the constrained folding data, plant pre-miRNA hairpins loops are significantly more stabilized than metazoans (**Fig. 3B**, **Supplementary table 2**), suggesting a distinguishing role of apical hairpin loop stability in metazoan miRNAs. While bulges are not significantly distinguished (p= 0.03), plant pre-miRNA internal loops are less stabilized than those of metazoa (**Fig. 3C-D**, **Supplementary table 2**). These observations are consistent with the role of loop-distal regions (49) and duplex stability (50) in modulating plant pre-miRNA processing.

**Figure 3.**
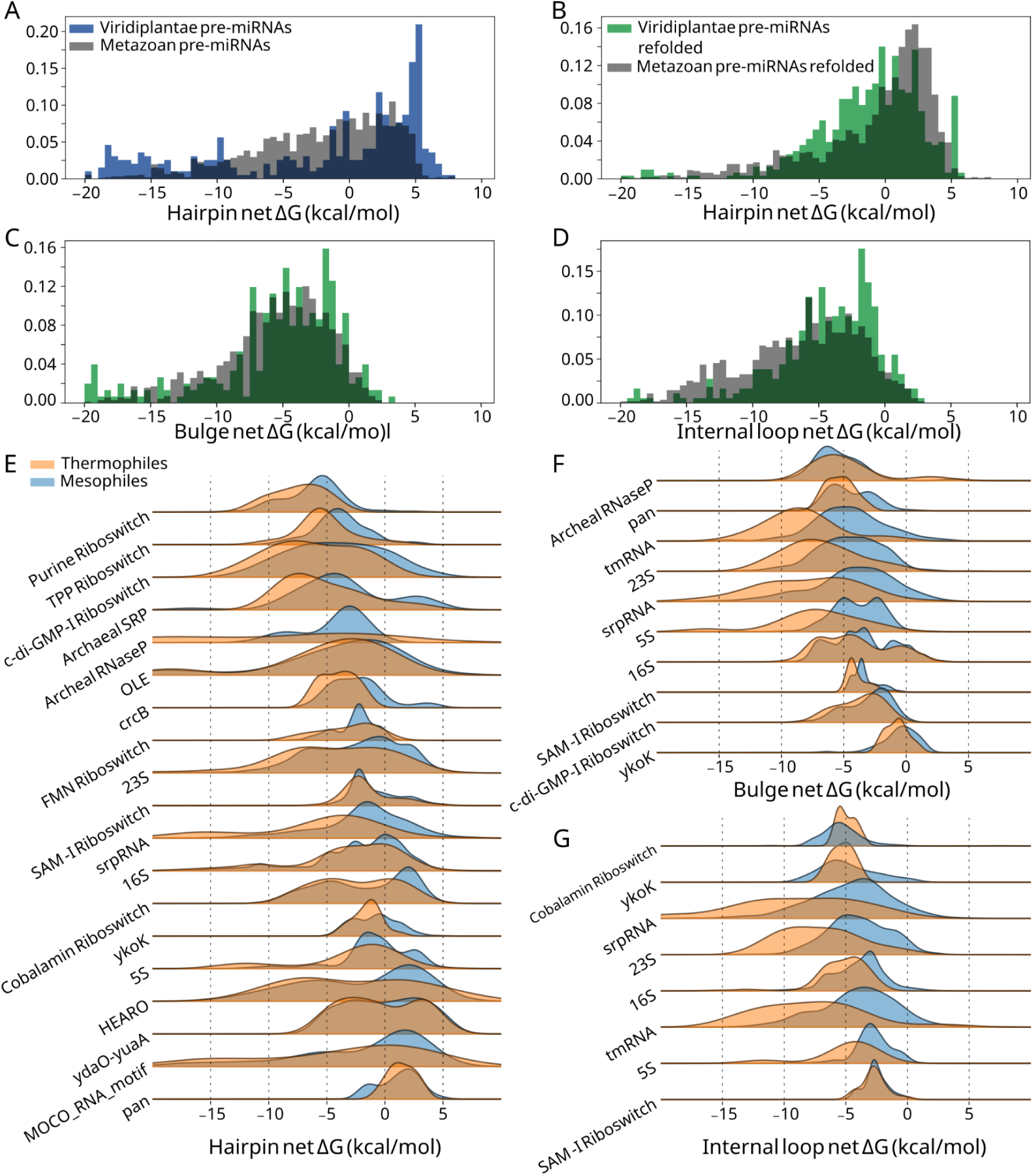
Differences in net ΔG by phylogeny for select RNA types. (A) plant pre-miRNA hairpin net ΔGs (green) and metazoan (grey) net ΔGs before folding with constraints and (B) after folding with constraints. (C) plant pre-miRNA bulge ΔGs (green) and metazoan (grey) net ΔGs after folding with constraints. (D) plant pre-miRNA internal loop ΔGs (green) and metazoan (grey) net ΔGs after folding with constraints. (E) Thermophilic (orange) hairpin net ΔGs and non-thermophilic net ΔGs (blue). (F) Thermophilic (orange) bulge net ΔGs and non-thermophilic net ΔGs (blue). (G) Thermophilic (orange) internal loop net ΔGs and non-thermophilic net ΔGs (blue).

Thermophilic species exhibit greater hairpin, bulge, and internal loop stability in ncRNAs relative to their mesophilic counterparts, in alignment with previous research showing increased GC content in tRNAs and rRNAs (39) (**Fig. 3E-G**). There are exceptions for which differences in local stability are either minute or absent, including regulatory elements such as the YkoK leader, ydaO, and the cobalamin riboswitch. The functional roles of participating substructures may explain the similarity in net ΔG for these structures. Interestingly, the S-adenosylmethionine riboswitch (SAM-I), shows strong agreement between thermophilic and non-thermophilic species for hairpins and internal loops (**Fig. 3E**, **3G**) yet a lower net ΔG exists for thermophilic SAM-I bulges (**Fig. 3F**). The only SAM-I bulge known to vary across phylogeny is a bulge adjacent to the SAM binding pocket, which is suggested to indirectly stabilize the pocket and promote the adoption of the binding-competent conformation (51). Deletion of this bulge is highly deleterious in *B. subtilis*, but not in *T. tengcongensis*, a thermophilic species, which may stabilize the binding pocket by other means such as a base triple (51). Thus, in an otherwise highly conserved riboswitch, a substructure has a lower net ΔG in thermophiles than it does in mesophiles. In light of these observations and in order to bypass confounding biological context, we devised an experiment to test LSC in rationally designed constructs.

### DMS reactivity of three sequence libraries demonstrates local stability compensation

To directly test LSC, we designed three RNA libraries that systematically varied local stabilities for assessment with dimethyl sulfate (DMS) chemical mapping. Each library used a template structure containing three bulges and one stem-loop, with randomized sequences in specific regions to generate variants with different local stabilities (**Fig. 4A**). This design allowed us to measure how stability changes affect both local and distal regions. To isolate the effects of local stability changes, we engineered constant-energy stems outside the variable regions as internal controls for our reactivity measurements.

**Figure 4.**
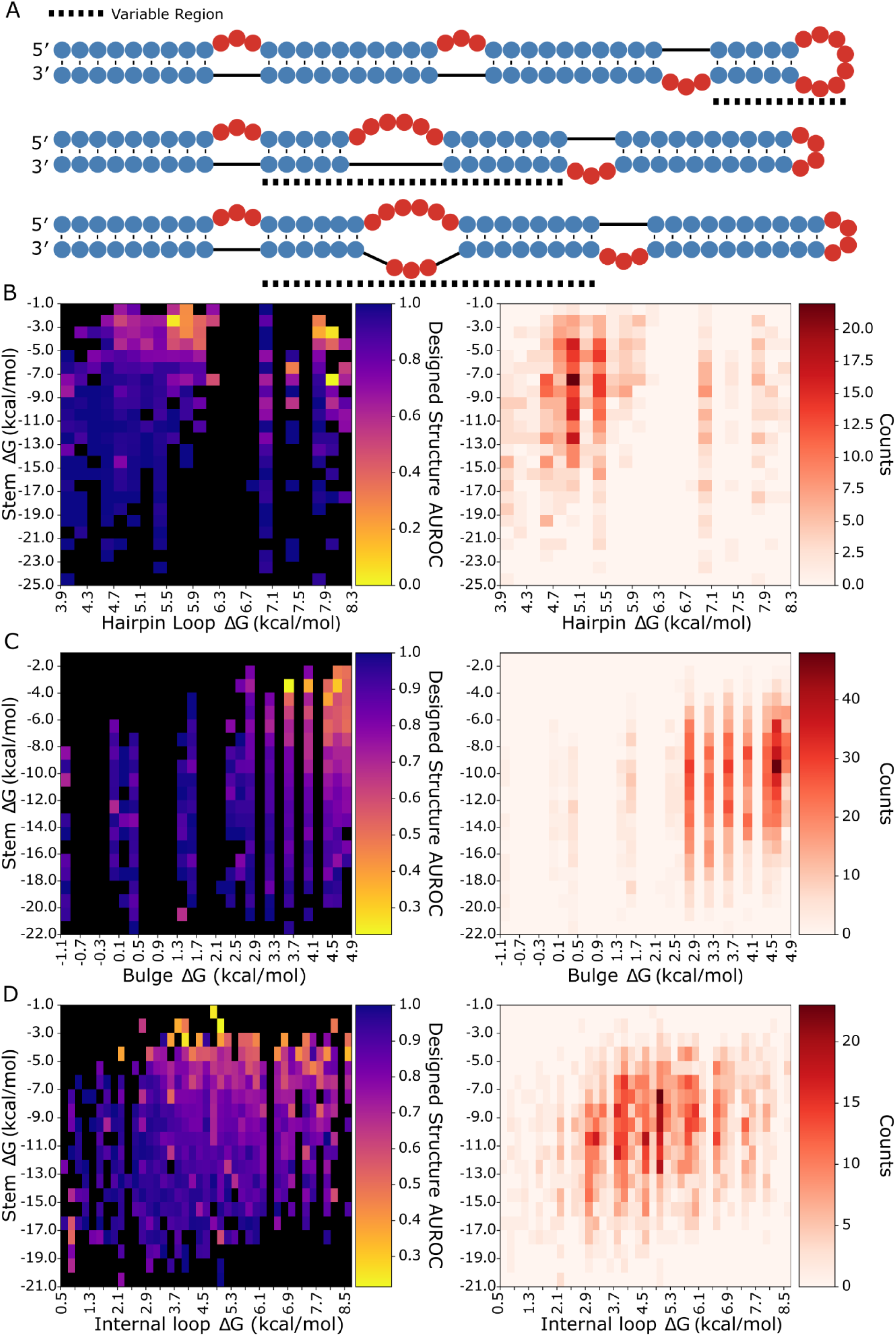
Library design and local structure AUROC heatmaps. (A) Designed structure examples for hairpins, bulges, internal loops, (top, middle, bottom respectively), dashed lines indicate the variable region. Average AUROC and structure count heatmaps for cells of (B) hairpin, (C) bulge, and (D) internal loop ΔG by adjacent stem ΔG, yellow cells show low AUROC or poor agreement of DMS data with the designed structure, while dark blue cells show high AUROC with strong agreement, and black cells contain no data. Count heatmaps show the number of RNAs contributing to each cell with darker red coloring indicating a higher number of RNAs.

Each library showed acceptable coverage of the distribution of net ΔGs in bpRNA-1m90 (**Supplementary Fig. 4A, 4C, 4E**). It is noted that the highest net ΔG bulges are underrepresented in the library when compared to their natural frequency in bpRNA-1m90. We used RNAfold (28) to predict each structure in the libraries and used the structure alignment approach bpRNA-align (52) to calculate a structural similarity score between each designed structure and its corresponding predicted structure, revealing that RNAfold correctness for the designed structures is unimpacted by loop ΔG (**Supplementary Fig. 4B, 4D, 4F**). The designed RNA structures were synthesized and DMS reactivity values were collected with a high-throughput sequencing approach (see methods).

To evaluate design success with a single metric, we used an established method to quantify the consistency between a given designed structure and the corresponding reactivity data (53). This was achieved by calculating the area under receiver operator curve (AUROC) of the DMS reactivity data against the designed target structure for each library (see methods). The values were binned by stem and loop free energy and are displayed in heatmaps (**Fig. 4B-D**). The heatmaps show lower average AUROC values for structures of higher loop ΔG and lower stem ΔG across each structure component type. In each heatmap, a roughly linear boundary exists past which local folding fidelity rapidly decreases, indicating that stem and loop energies both contribute to local folding fidelity. Furthermore, AUROC correlations are strongest with net ΔG, although stem ΔG correlates more than loop ΔG (Supplementary Table 4A). It is noted that a number of low stem energy, low loop energy bulges and internal loops have poor AUROC values. As a control for the non-local structure, the AUROC scores for the remaining region are unaffected by the net ΔG of the local, variable region (**Supplementary Fig. 5**).

### Loop stabilization primarily impacts adjacent stem reactivities

We investigated which components of the structures in the library were changing in low AUROC structures (**Fig. 5**). For both local and distal helices, we calculated the average stem reactivity, which is a more direct readout of helix formation than local AUROC. Locally, average stem reactivity correlates well with stem-loop net ΔG and shows a strong positive trend. Correlation R-squared values were 0.458, 0.259, 0.318 for hairpin loops, bulges and internal loops respectively (**Fig. 5B, 5F, 5J**). These correlations are stronger than loop or stem ΔGs alone (Supplementary Table 4B). This indicates that stems closing less stabilized loops either adopt partially folded states or completely unfold. Distal stem reactivity shows no correlation with net ΔG (**Fig. 5C**, **5K**), except for a weaker but slightly positive trend in bulges (**Fig. 5G**). These data show that RNA stem loop instabilities have predominantly local effects.

**Figure 5.**
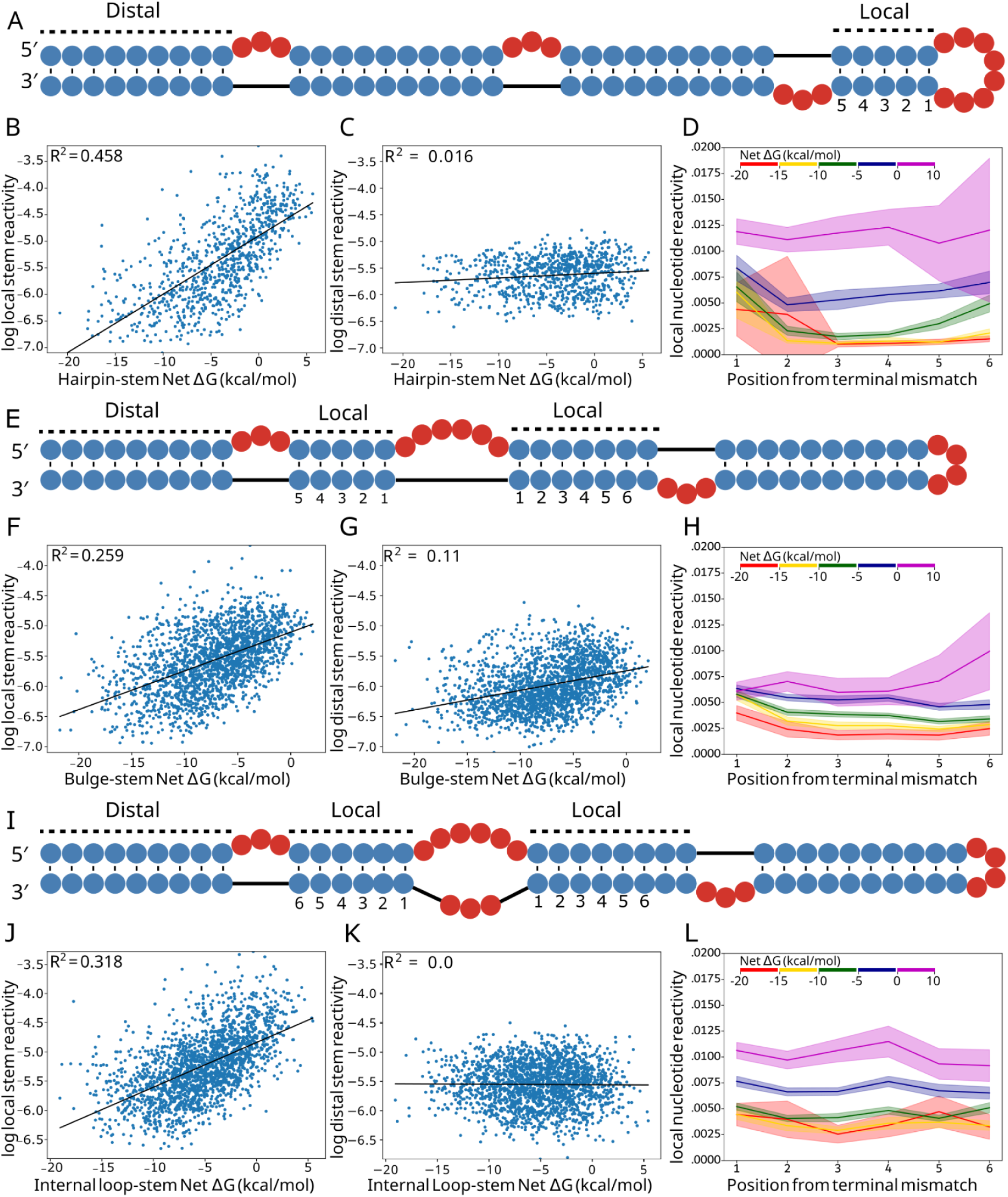
Local vs distal stem reactivity correlations. (A) Example structure from the hairpin library with adjacent and distal stems to the hairpin loop labeled and local stem base pair indices labeled. (B) Linear regression between hairpin net ΔG and log average local stem reactivity, compared to average distal (C) stem reactivity. (D) Average net ΔG bin reactivity per position away from the hairpin loop. (E) Example structure from the bulge library with adjacent and distal stems to the bulge labeled and local stem base pair indices labeled. (F) Linear regression between bulge net ΔG and log average local stem reactivity, compared to average distal (G) stem reactivity. (H) Average net ΔG bin reactivity per position away from the bulge, where an index is present on both stems, the average is used. (I) Example structure from the internal loop library with adjacent and distal stems to the internal loop labeled and local stem base pair indices labeled. (J) Linear regression between internal loop net ΔG and log average local stem reactivity, compared to average distal (K) stem reactivity. (L) Average net ΔG bin reactivity per position away from the internal loop, where an index is present on both stems, the average is used.

We next sought to investigate how insufficiently compensated stems were impacted at the residue level. We calculated the average DMS reactivity of each base pair succeeding the loop for stems belonging to five different net ΔG bins. Hairpins, bulges, and internal loop averages and 95% confidence intervals are plotted (**Fig. 5D, 5H, 5L**) up to position 6, after which too few data points exist for the higher energy bins. For bulges and internal loops, values for positions present on both stems were averaged. In agreement with average stem reactivity results, hairpin loop, bulge, and internal loop per-nucleotide reactivity distributions increase monotonically with each ΔG bin. Stem reactivity increases more for the closing base pair (position 1) than for the subsequent pairs. Stems closing hairpin loops show this behavior more strongly and for lower free energy bins than those closing bulges and internal loops, possibly due to strain involved in loop closure for hairpins. Bins of higher net ΔG experience dramatic shifts in reactivity for the middle of the stem (positions 2-5), and for all loop types, the highest energy bin (ΔG > 0 kcal/mol) shows reactivity consistent with unpaired nucleotides for the entire stem. Importantly, the net ΔG range between -5 kcal/mol and 0 kcal/mol shows reactivity values that are intermediate between paired reactivities and unpaired reactivities. This intermediate category is the most represented in bpRNA-1m90 histograms, while fewer substructures in the database belong to the high energy bin (0 kcal/mol to 10 kcal/mol), and still fewer belong to the lowest energy bin (-20 kcal/mol to -15 kcal/mol) (**Supplementary Fig. 4A, 4C, 4E**).

To investigate other reactivity changes possibly responsible for the observed AUROC values, average loop reactivities were evaluated across stem-loop net ΔGs (**Supplementary Fig. 6A-C**), revealing weak trends towards lower reactivity with respect to stem-loop net ΔG. These trends are similar in strength and magnitude across loop types, and the trends for local loop reactivity are marginally stronger than those of the distal loop. It is expected that loop reactivities can significantly decrease when a bulge or internal loop shifts (**Supplementary Fig. 7B**). While these weak negative trends in loop reactivity do not compare to the correlations found between substructure net ΔG and stem reactivity, they partially contribute to local AUROC values. Thus, loop residues are not primarily responsible for changes in AUROC with increasing net ΔG, rather, it is the degree to which stems compensate for loop closure that produces the observed changes in local AUROC.

### Local substructure net free energies are correlated to local folding fidelity and fit the Hill equation

We next sought to define a relationship between LSC and local folding fidelity by comparing net ΔGs with the AUROC values for each structure type such that a given net ΔG estimates the resulting folding behavior. Highlighted in the hairpin loop scatterplot (**Fig. 6A**) are three examples showing high, medium and low agreement with the designed structure (**Fig. 6B**). Structures with low AUROC (i.e. 0.25) contain many bases that were designed to be paired yet have higher reactivity than those of the adjacent loop.

**Figure 6.**
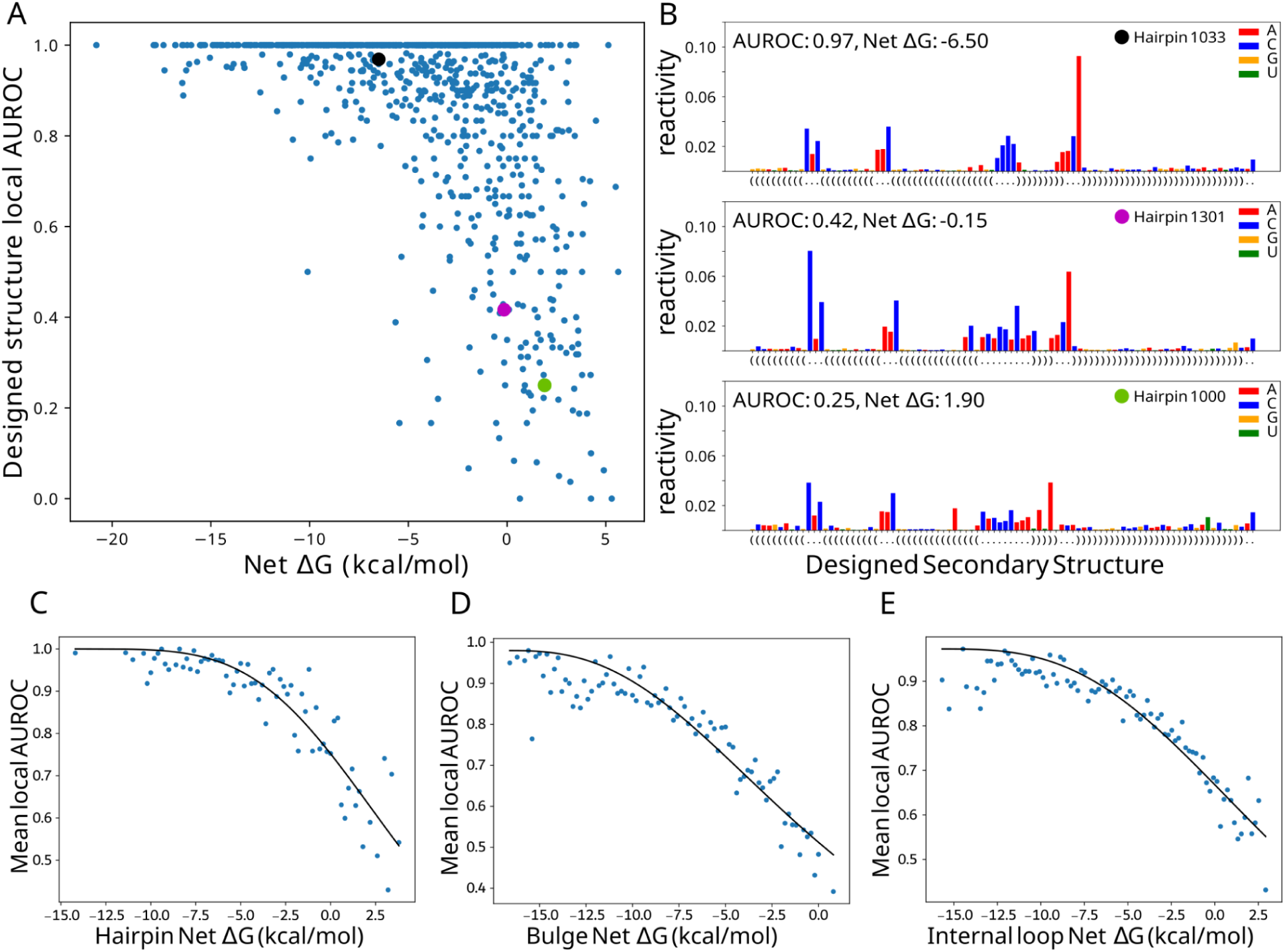
Local structure fidelity curve fitting. (A) Hairpin net ΔG vs average local AUROC, points indicated by black, magenta, and cyan are shown in (B), raw reactivity values color-coded by nucleotide for three example designed hairpins, the boxed region is involved in AUROC calculation. (C,D,E) Hill equation curves fit to net ΔG bins (0.2 kcal/mol) versus the mean AUROC for each bin.

In order to reduce noise associated with the DMS reactivity data and the AUROC calculation, AUROC data was averaged across bins of 0.2 kcal/mol net ΔG. The observed local AUROC asymptotically approaches 1.0 for decreasing net ΔG, suggesting a two-state model that includes a state that folds according to the design and a divergent folding state at low AUROC values. This observation of a two-state model with saturation led us to fit the averages to the Hill equation (**Fig. 6C-E**). Hairpin, bulge, and internal loop AUROC data cross an AUROC of 0.9, which represents minor DMS data disagreement, at -4, -9, and -6 kcal/mol, respectively (**Fig. 6C-E**). These thresholds show the least stability needed to fold with reasonable consistency. Hairpin AUROC data show a steeper curve (larger Hill coefficient) (**Supplementary Table 5**) and a clearer fit line, indicating that hairpin loops are less tolerant of reductions in stability, compared to bulges and internal loops. Relative to hairpins and internal loops, bulges require a greater degree of stabilization, as indicated by the more negatively shifted curve. The consistency of the fit to the Hill equation suggests that net ΔG is, on average, an effective predictor of local folding fidelity.

## Discussion

Here we demonstrate that local stability compensation, wherein loops are stabilized by their adjacent helices, can be observed within an RNA meta-database and in DMS reactivity data collected for three structure libraries. The inherent diversity of structures and functions throughout RNA families leads to varying levels of LSC, and the clearest trends are observed for bulges and internal loops, while a wider range of net energies exists for hairpins. By studying three libraries of template-based RNA designs, we show a degree of independence of local stability on the remainder of the structure, and reveal how the net free energy of substructures influence the folding of the involved stem.

Differences in local substructure stability exist between RNAs of different form and function. Bulges, which showed the strongest correlation between stem and loop-free energy, had the least variance in net free energy by RNA type, while the opposite was observed for hairpins. The structural and functional contexts of different RNA types also play a role in stability variation as the Turner parameters do not capture the free energy contributions of ligand binding, non Watson-Crick-Franklin base pairs, and tertiary interactions—an important limitation of our study. Additionally, we averaged the free energies of flanking stems of bulges and internal loops, which is an efficient way to define a “local” substructure; however, this approach overlooks stem symmetry and obfuscates isolating one substructure from another. A more refined model for determining locality should be pursued to address this limitation.

Many RNA families show patterns that are consistent with LSC. The tRNA T, D, and anticodon loops were clustered into distinct stem and loop free energy regions (**Fig. 2F)**. While the anticodon loop is typically larger, possibly to accommodate other interactions such as with the codon, it also shows a more stable associated stem, consistent with the LSC hypothesis. C4 antisense RNAs are characterized by two hairpin loops consistent with LSC: H2 is larger, and has an adjacent stem with high GC content, while H1 is a highly stable tetraloop and is closed by a stem with low GC content (**Fig. 2B**). Lastly, this observation may be applied to unknown structures. For example, the TwoAYGGAY motif has two identical hairpin loops which show distinct patterns of net free energy (**Fig. 2D**), suggesting different structural contexts which may be of interest for future research.

Such functionally relevant LSC patterns include taxonomy-specific distinctions. For example, metazoan microRNA hairpin loops exhibited an enrichment of the highest net ΔGs, which may be consistent with functions of the apical loop. DGCR8 (54) and hemin (55) interact at the UGU motif in the apical loop, as well as a peptide (56) designed to prevent miRNA maturation. Additionally, the instability of the stem adjacent to the hairpin loop may facilitate the adoption of the local helical distortion favorable for interactions with the dsRBD of DICER (57). In contrast, plant pre-miRNAs, which do not involve the hairpin loop for maturation, do not exhibit high energy hairpin loops, but instead show an enrichment of high energy internal loops. Taken together, these observations suggest that substructure net free energy is linked to biological context. Furthermore, patterns in net free energy may provide clues to uncover functions and highlight biological differences that could drive future investigations.

The designed structure libraries revealed contributions of stem and loop free energy to folding fidelity, which solidifies net ΔG as an appropriate metric for LSC. The primary factor underlying a substructure’s folding fidelity was the reactivity of the involved stem or stems. Comparing these reactivities against those of distant stems revealed that LSC impacts the local stem much more than distal stem, suggesting that local substructures maintain a degree of independence or modularity on the thermodynamic level. Notably, the distal reactivities for bulges showed a detectable trend unlike the distal reactivities for hairpins and internal loops, which could be related to the underrepresentation of high energy bulges (**Supplementary Fig. 4**). Loops were found to decrease in reactivity with respect to net ΔG, possibly due to the influence of non-canonical interactions or artifacts of successive reactive nucleotides (43).

The relationship between net ΔG and folding fidelity is half sigmoidal, with substructure free energies around -20 to -10 kcal/mol plateauing at an AUROC of 1.0 (**Fig. 6C,D,E**). Such over-stabilized structures are very rare in naturally occurring structures, consistent with the idea of an “energy budget”, wherein G-C base pairs have a tendency to not be overused because of the diminishing returns as indicated by the plateauing trend. In contrast, intermediate net ΔG values (-10 to 0 kcal/mol) are abundant in naturally occurring structures. The reactivity data (**Fig. 5D,H,L**) shows intermediate reactivity across these substructures and may be explained by partial folding, transient folding, or likely an ensemble of folding states (58–60), which are implicated in RNA function. Under-stabilized structures with a net ΔG greater than 0 kcal/mol are infrequent, and could possibly represent substructures that require instability or are stabilized by external interactions (e.g. microRNA hairpin loops interacting with hemin and DGCR8) or tertiary interactions. Tertiary motifs involve non-local interactions and were not included in the study, however, it is likely that the local stability of participating substructures may influence tertiary motifs or their formation. Investigating the relationship between local stability compensation and tertiary contact formation would be a logical next step for future research.

In MFE RNA structure prediction, randomized sequences can fold with a similar MFE compared to the original sequence (61), and MFE is not distinguished between known structured RNA and random sequences (62). Additionally, since the number of possible helices increases exponentially with length (61), it is clear that additional constraints are needed to improve thermodynamic structure prediction. Our results suggest that local free energy balance in addition to global minimum free energy will improve thermodynamic approaches for structure prediction as well as the inverse folding of RNA.

Here, we describe a thermodynamic locality that occurs fundamentally in RNA structure and has widespread implications for the rational design of RNA, where both sequence and structure are engineered to achieve a desired outcome. According to the presented relationships between LSC and local folding behavior, an RNA engineered from well-folding substructures may also be expected to fold well, and an otherwise stable RNA may include intermediate to under-stabilized substructures, since the effects of this instability on the remainder of the structure are mostly insignificant (**Fig. 5, Supplementary Fig. 5**). Further work with RNA libraries and high-throughput chemical mapping could reveal similar relationships in multiloops and other RNA motifs, as well as uncover the role of LSC in small binding sites. With further characterization, LSC could serve as a guide for the design of future biotechnology including molecular machines composed of structured RNAs.

## Supporting information

Sequences.xlsx

## Data availability

All data, materials, and software used in this study are available. Unprocessed FASTQ files have been deposited to the Sequence Read Archive (SRA) under the accession SRP548417. All other data is available on Fig Share (https://doi.org/10.6084/m9.figshare.27281064.v1). All original code used in this study is available at Github (https://github.com/cornwero/LocalStabilityCompensation) and Zenodo (https://doi.org/10.5281/zenodo.14252189) and full datasets are available at figshare (https://figshare.com/ndownloader/files/51111164).

## Author Contributions

DAH and JDY conceived this study. RN performed the experiments. RCA performed the analysis of the bpRNA-1m meta database and chemical mapping data with assistance from DAH and JDY. MTH wrote the original code to parse and calculate free energies for RNA substructures. DAH, JDY and RCA wrote the paper with assistance from RN.

## Acknowledgements

The authors would like to acknowledge the Center for Quantitative Life Sciences (CQLS) at Oregon State University for maintaining the server infrastructures on which the analysis was performed. This work was completed utilizing the Holland Computing Center of the University of Nebraska, which receives support from the UNL Office of Research and Economic Development, and the Nebraska Research Initiative. Additionally, the authors would like to acknowledge Aidan B. Estelle and Chandler Peterson for their early conceptual contributions to the project.

