## Supplemental for "Analysis of natural structures and chemical mapping data reveals local stability compensation in RNA"

### Supplementary Information

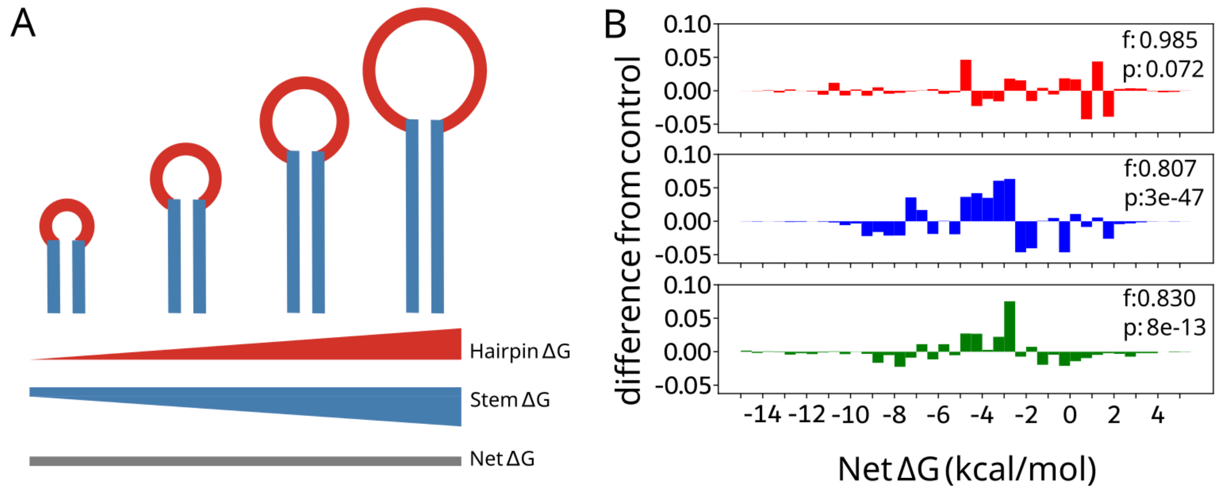

**Supplementary Figure 1. Local stability compensation against an in-structure rotation control.**(A), Schematic demonstrating local stability compensation, which proposes that larger hairpins are closed by larger stems to maintain the net free energy. (B) Diagram explaining stem swapping in the rotation control (C) The difference in net ΔG distribution between bpRNA-1m90 refolded structures versus rotated structures for hairpins (red)(p=0.072), bulges (blue)(p=3e-47), and internal loops (green)(p=8e-13).

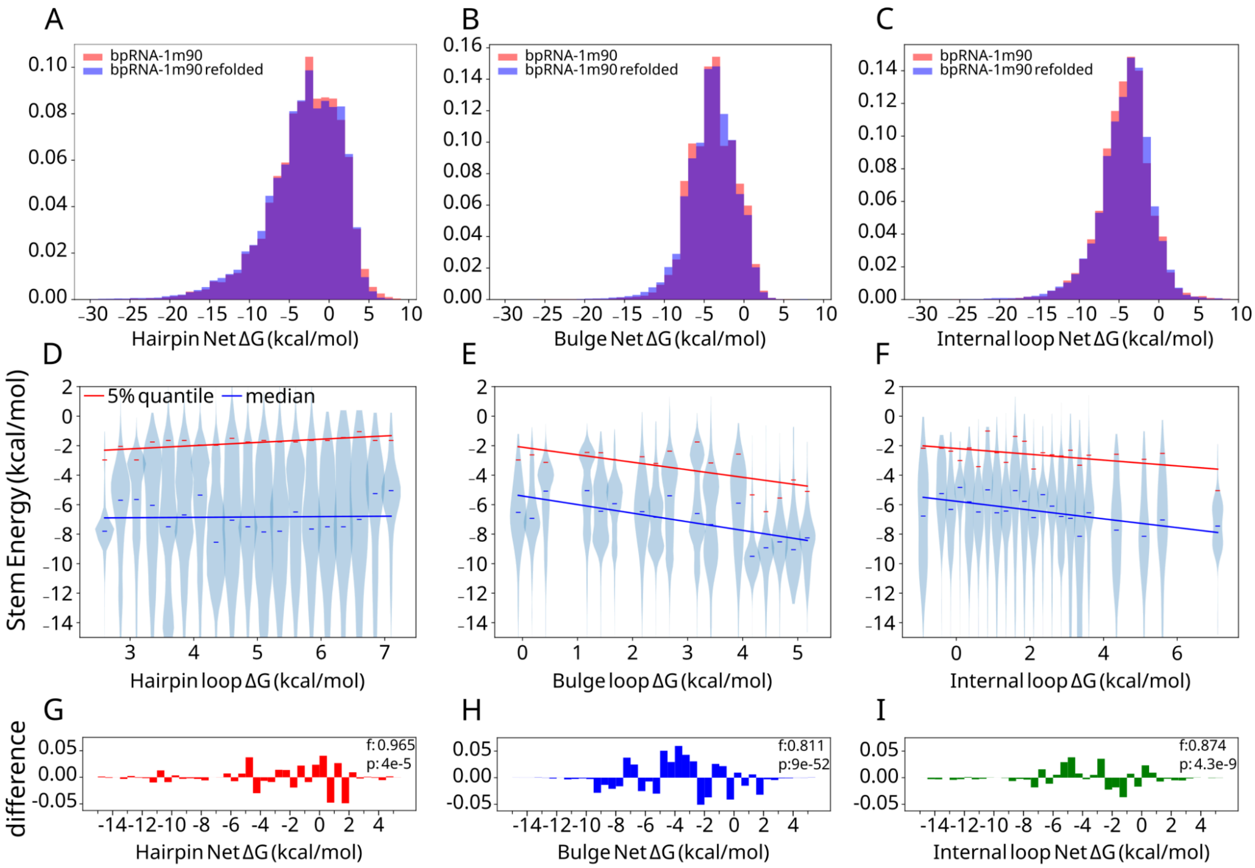

**Supplementary Figure 2. Comparing refolded data with bpRNA-1m90.** (A) Hairpin (B) bulge and (C) internal loop net ΔG densities for bpRNA-1m90 and bpRNA-1m90 including refolded structures. (D) Hairpin, (E) bulge, and (F) internal loop ΔGs are binned and the corresponding stem ΔGs are plotted as violins with linear regressions drawn for the median (blue) and 5% quantile (red). (G) The difference in net ΔG distribution between bpRNA-1m90 structures versus rotated structures for hairpins (red)(p=4e-5), (H) bulges (blue)(p=9e-52), and (I) internal loops (green)(p=4.3e-9).

| | Hairpins | Bulges | Internal Loops |
|---|---|---|---|
| 5% quantile refolded | 0.370 | 0.412 | 0.456 |
| median refolded | 0.401 | 0.647 | 0.631 |
| 5% quantile | 0.451 | 0.418 | 0.245 |
| median | 0.001 | 0.506 | 0.404 |

**Supplementary Table 1. Correlations of stem ΔG median and 5% quantile in refolded data and bpRNA-1m90.**

| | Hairpins | Bulges | Internal Loops |
|---|---|---|---|
| Mean plotted RNA classes | -2.66 | -4.29 | -3.85 |
| Stdev plotted RNA classes | 1.74 | 0.75 | 1.33 |
| Mean all subclasses | -2.64 | -4.90 | -4.59 |
| Stdev all subclasses | 5.44 | 3.35 | 3.77 |

**Supplementary Table 2. Average and standard deviation of RNA category medians.** Plotted RNA classes include only the major RNA classes included in figure 1.

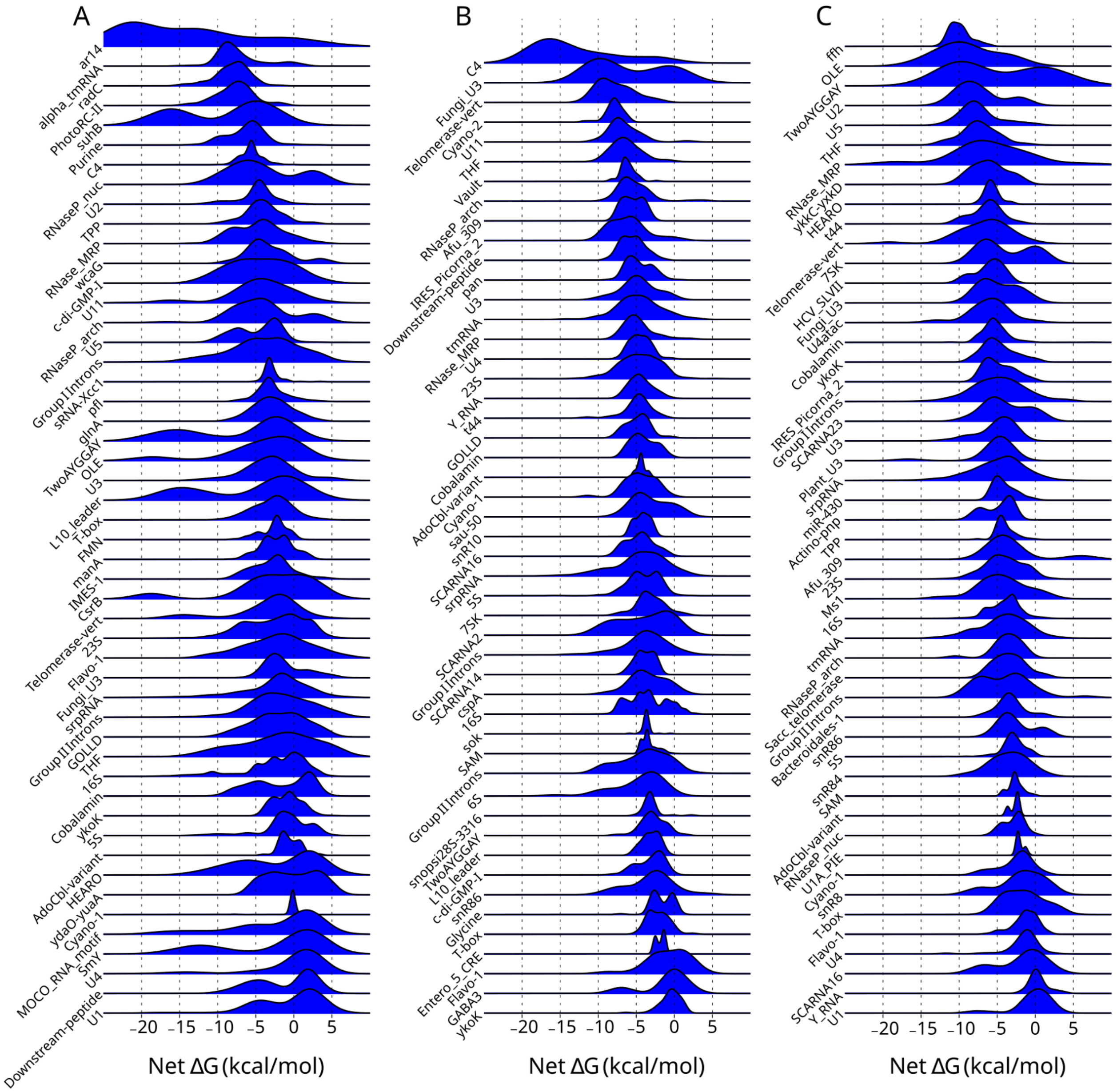

**Supplementary Figure 3. Net ΔG variation by RNA subtype in bpRNA-1m90.** Subtypes sorted by median net ΔG and the corresponding net ΔG distributions are plotted for (A) hairpins, (B) bulges, and (C) internal loops.

| | Hairpins (pre-refold) | Hairpins | Bulges | Internal loops |
|---|---|---|---|---|
| KS statistic | 0.207 | 0.166 | 0.088 | 0.195 |
| p value | 0.000 | 0.000 | 0.035 | 0.000 |

**Supplementary Table 3. plant pre-miRNA net ΔGs are distinct from those of other species.**

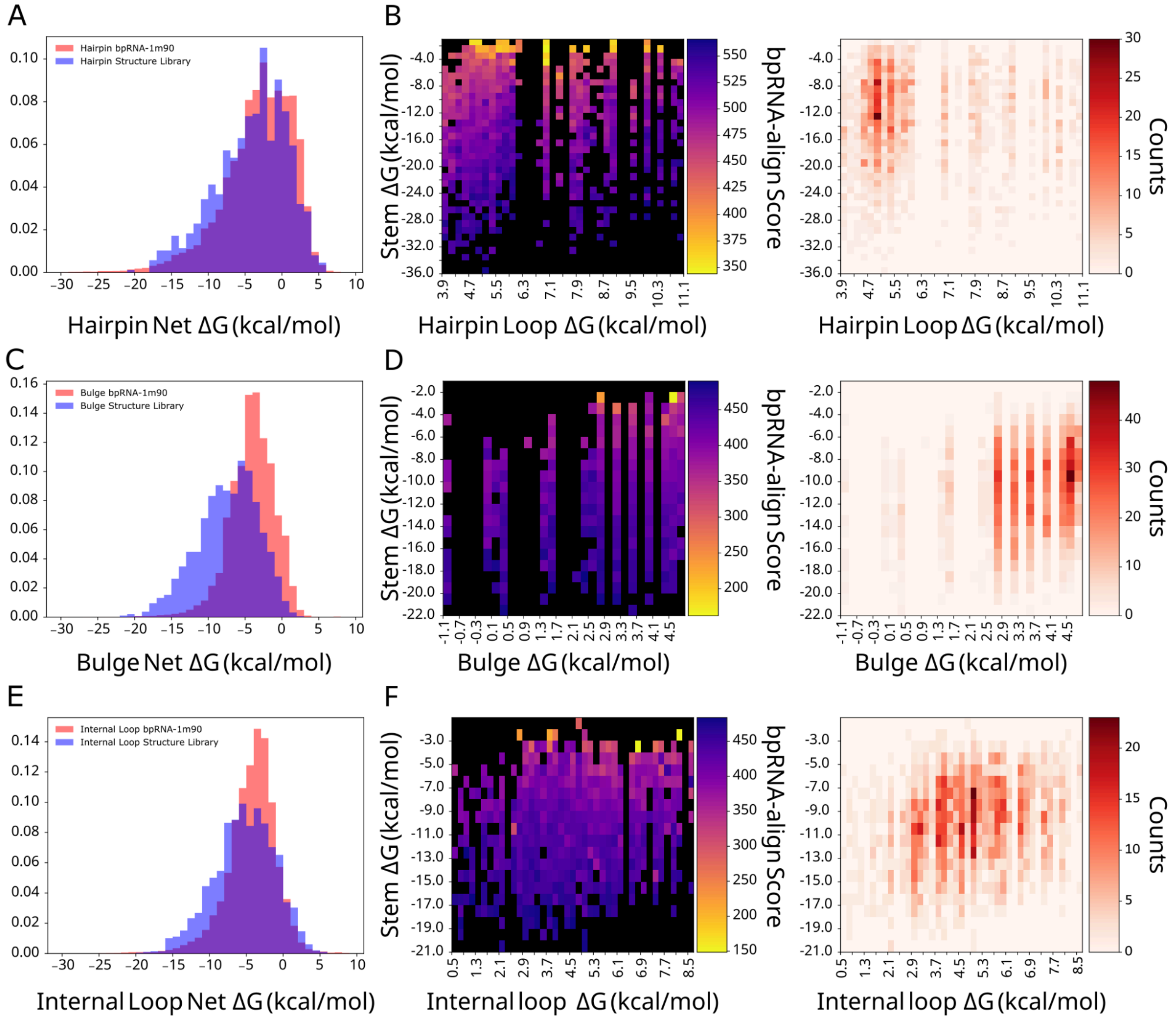

**Supplementary Figure 4. Library validation and RNAfold structural similarity heatmaps**. (A) Hairpin library net ΔG overlap with refolded RNAs from bpRNA-1m90, (B) Structure alignment score heatmaps for library structure hairpin and stem ΔG, sample sizes for each cell. (C) Bulge library net ΔG overlap with bpRNA-1m90, (D) Structure alignment score heatmaps for library structure bulge and stem ΔG, sample sizes for each cell. (E) Internal loop library net ΔG overlap with bpRNA-1m90, (F) Structure alignment score heatmaps for library structure internal loop and stem ΔG, sample sizes for each cell. Heatmap cells colored yellow have poor alignment while cells colored blue have good alignment.

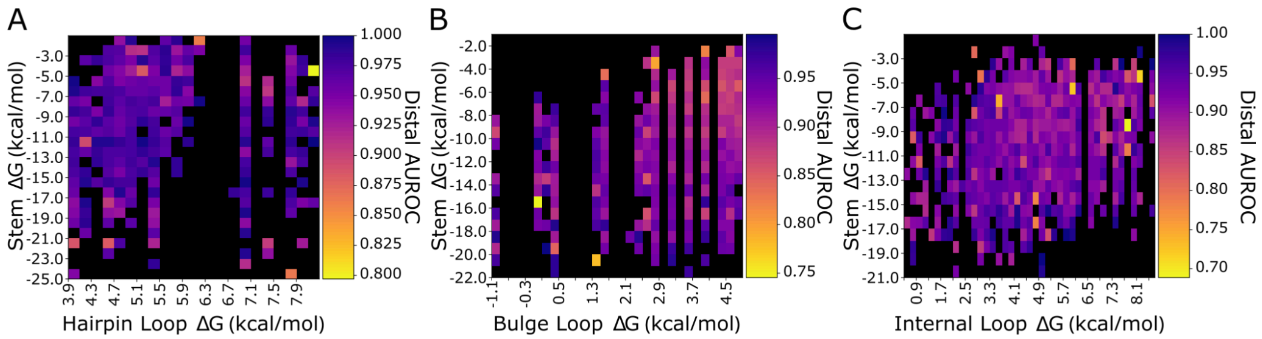

**Supplementary Figure 5. Non-local AUROC is not modulated by local stem or loop ΔG**. AUROC heatmaps for the distal structures of (A) Hairpin loops, (B) bulges, and (C) internal loops.

| A) Local AUROC | Loop ΔG $R^2$ | p | Stem ΔG $R^2$ | p | Net ΔG $R^2$ | p |
|---|---|---|---|---|---|---|
| Hairpins | 0.055 | 3e-12 | 0.226 | 3e-50 | 0.260 | 7e-59 |
| Bulges | 0.068 | 5e-32 | 0.187 | 5e-91 | 0.211 | 1e-103 |
| Internal Loops | 0.069 | 1e-32 | 0.179 | 7e-87 | 0.206 | 1e-101 |

| B) Local stem reactivity | Loop ΔG $R^2$ | p | Stem ΔG $R^2$ | p | Net ΔG $R^2$ | p |
|---|---|---|---|---|---|---|
| Hairpins | 0.045 | 3e-10 | 0.426 | 7e-107 | 0.458 | 9e-118 |
| Bulges | 0.104 | 3e-49 | 0.219 | 4e-108 | 0.258 | 4e-130 |
| Internal Loops | 0.083 | 3e-39 | 0.295 | 7e-153 | 0.318 | 2e-167 |

**Supplementary Table 4. Net ΔG is best correlated with folding metrics.** (A) Local AUROC correlation coefficients corresponding to loop, stem, and net free energy. (B) Local stem reactivity correlation coefficients corresponding to loop, stem, and net free energy.

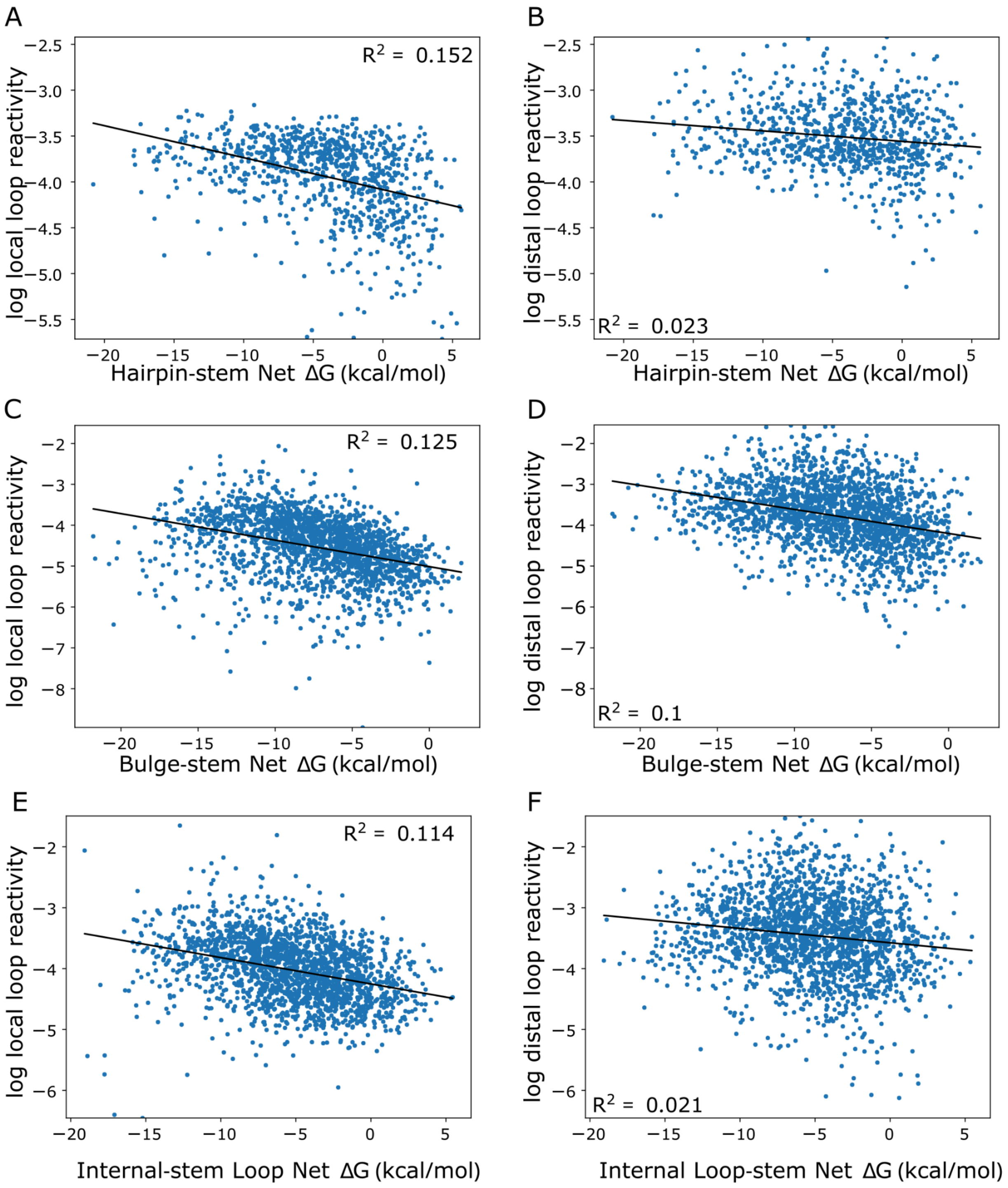

**Supplementary Figure 6. Loop reactivities are invariant to loop net ΔGs.** Hairpin-stem net ΔG versus local (A) and distal (B) average loop reactivities. Bulge-stem net ΔG versus local (C) and distal (D) average loop reactivities. Internal loop-stem net ΔG versus local (E) and distal (F) average loop reactivities.

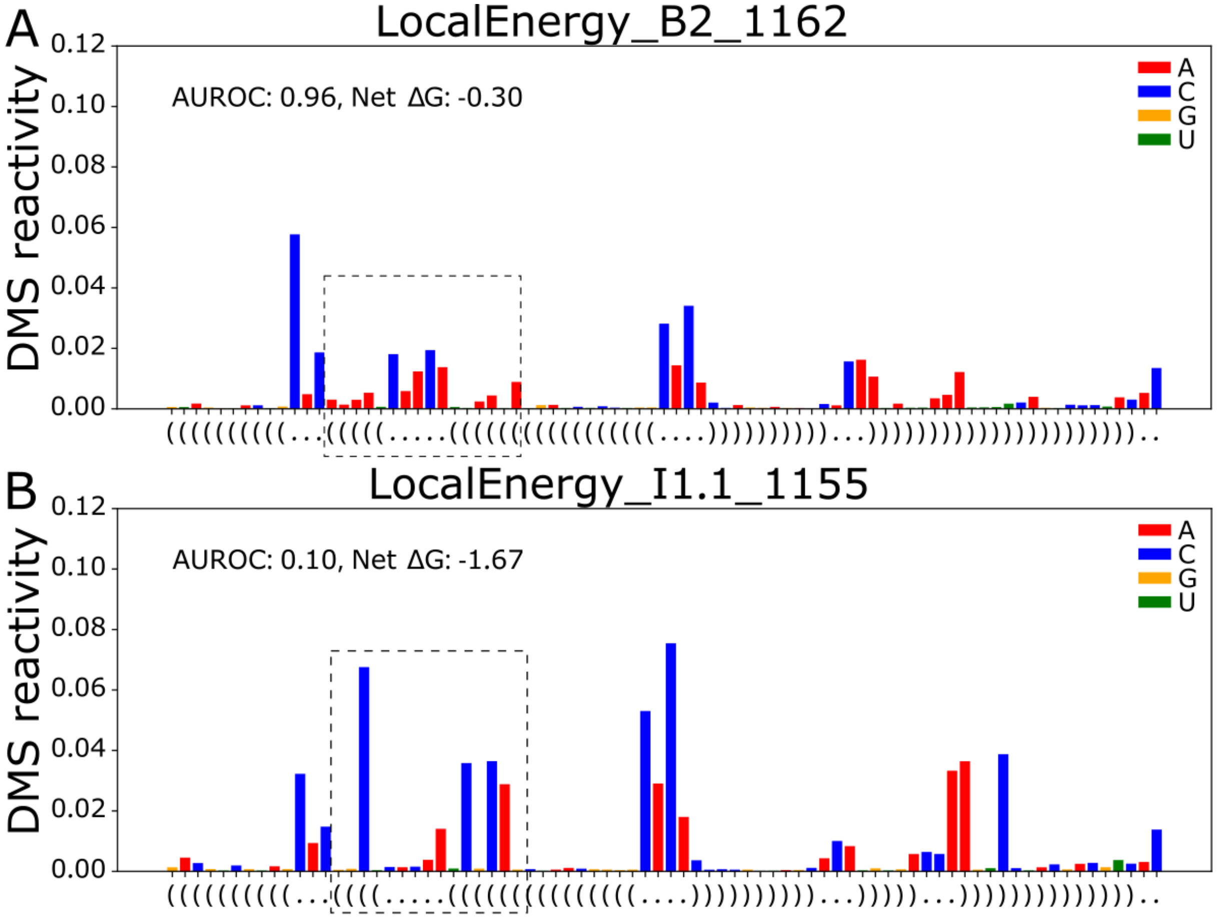

**Supplementary Figure 7. Designed structure reactivity examples.** (A) example structure from the bulge library with the bulge and adjacent stems highlighted. (B) an example internal loop showing loop shifting in the highlighted region.

| | K | n | A |
|---|---|---|---|
| Hairpin loops | 18.6 | 4.10 | 1. |
| Bulges | 17.4 | 2.58 | 1. |
| Internal loops | 20.7 | 2.95 | 1. |

**Supplementary Table 5.** Fit parameters for curve fitting to the Hill Equation.
